# Environmental context as a key driver of *Pseudomonas*’ biocontrol activity against *Salmonella*

**DOI:** 10.1101/2025.06.23.661019

**Authors:** Noemie Vimont, Sarah Bastkowski, George M. Savva, Samuel J. Bloomfield, Alison E. Mather, Mark A. Webber, Eleftheria Trampari

## Abstract

*Salmonella* poses a significant threat to food security, with frequent outbreaks reported worldwide. A large percentage of these outbreaks are associated with fresh produce intended for raw consumption. Plant microbiomes harbour diverse microbial communities, including commensal microbes such as *Pseudomonas* that sometimes exhibit biocontrol activity against plant pathogens. However, little is known about whether *Pseudomonas* strains can effectively suppress foodborne pathogens and the mechanisms they employ. In this study, we identified and characterised food derived *Pseudomonas* isolates capable of inhibiting *Salmonella* growth *in vitro* and *in planta*. The identified isolates were active against a range of *Salmonella* serovars, and additionally *E. coli* isolates derived from food. We demonstrated that dynamics of the interaction between *Pseudomonas* and *Salmonella* are environment and application dependant. To uncover the mechanisms *Pseudomonas* employs to suppress *Salmonella*, we used transcriptomics coupled with pathway analysis in the two different settings. We showed that *Pseudomonas* metabolism undergoes significant environment-specific changes in the presence of *Salmonella*, implicating different pathways responsible for the control of the pathogen in the two different settings. Our results highlight the plasticity of *Pseudomonas* metabolism in response to *Salmonella* in two distinct environments and provide evidence that *Pseudomonas* biocontrol activity is multifactorial and environment dependent.

## Introduction

*Salmonella* is a major foodborne pathogen responsible for numerous outbreaks (*1*). *Salmonella’s* ability to use alternative hosts such as plants (*2*) to survive and proliferate in diverse environments, renders fresh produce a significant source of reported outbreaks associated with *Salmonella* (*3*). Previous work has demonstrated how *Salmonella* employs specific mechanisms to colonise plant tissues, leading to increased challenges with infection control and disinfection of fresh produce (*4*,*5*).

In nature, bacteria live in multi-species ecosystems, communities shaped by complex dynamics. Competitive interactions drive microbial community structure, diversity and function (*6*). Microbes employ an arsenal of tools to compete for space and resources (*7*). Organisms such as *Pseudomonas* have been well characterised and studied for their “biocontrol activity”, the ability to control the growth of pathogens (*8*, *9*). *Pseudomonas fluorescens* (*P. fluorescens*) uses distinct suppression mechanisms against plant pathogens and other competing microbes (*10–12*).

One key mechanism is the production of antimicrobial compounds. *P. fluorescens* produces secondary metabolites with antimicrobial properties, including 2,4-diacetylphloroglucinol (DAPG), pyoluteorin, pyrrolnitrin, and hydrogen cyanide (*13*), which exhibit broad-spectrum activity against various fungi and bacteria. Another mechanism is the production of siderophores, particularly pyoverdine, which allows *P. fluorescens* to scavenge iron from the environment (*14*), thereby depriving potential pathogens of iron and inhibiting their growth. This competitive iron acquisition plays a significant role in the biocontrol activity of *Pseudomonas* against plant pathogens (*15*,*16*).

Furthermore, the metabolic versatility of *P. fluorescens* allows it to use a wide range of organic compounds as carbon and energy sources, giving it an advantage in nutrient-limited environments (*17*). This ability to rapidly colonise and efficiently use available nutrients contributes to its effectiveness in outcompeting pathogens.

*P. fluorescens* is not only relevant in fresh produce and plant ecosystems, but also in other food chain related environments. It is one of the most prevalent species in food processing environments, making it an integral part of these ecosystems (*18*). Understanding the principles governing antagonistic interactions involving *Pseudomonas* and human pathogens in different environments is key to promoting our understanding of community-level dynamics, which we can in turn exploit to prevent human pathogens from establishing and disseminating in food and food processing environments.

This study explores the interplay between *P. fluorescens* and *Salmonella enterica* in two distinct environments: *in vitro* and *in planta*. Employing high throughput phenotyping techniques, competition assays, transcriptomics and pathway enrichment analysis, we showed that *P. fluorescens* effectively competes against *Salmonella* in both environments tested. We showed there is no universal mechanism employed by *Pseudomonas* to compete against *Salmonella.* Instead, *P. fluorescens* undergoes significant metabolic shifts which are likely to determine the mechanisms used to control *Salmonella* in an environment specific manner.

## Results

### *Pseudomonas* efficiently inhibits the growth of *Salmonella in vitro*

We assessed a panel of 54 isolates belonging to the *P. fluorescens* complex that had been previously isolated from food products (*18*) for their ability to inhibit *Salmonella enterica* Typhimurium 14028S. We selected strains based on their ability to inhibit *Salmonella* using a high-throughput competition assay on agar.

Our analysis revealed distinct clustering particularly for strains exhibiting high activity against *Salmonella*, with these isolates grouping closely together in the phylogenetic tree (Figure 1a). Our downstream investigations focused on a strain that consistently demonstrated Highly Potent Biocontrol Activity against *Salmonella* (BA_HP_), in all assays and environments tested as described below. The strain, which was originally isolated from pork, was whole genome sequenced and classified as *Pseudomonas lactis* according to the Genome Taxonomy Database, which places it within the broader *P. fluorescens* complex (Figure 1a).

**Figure 1:**
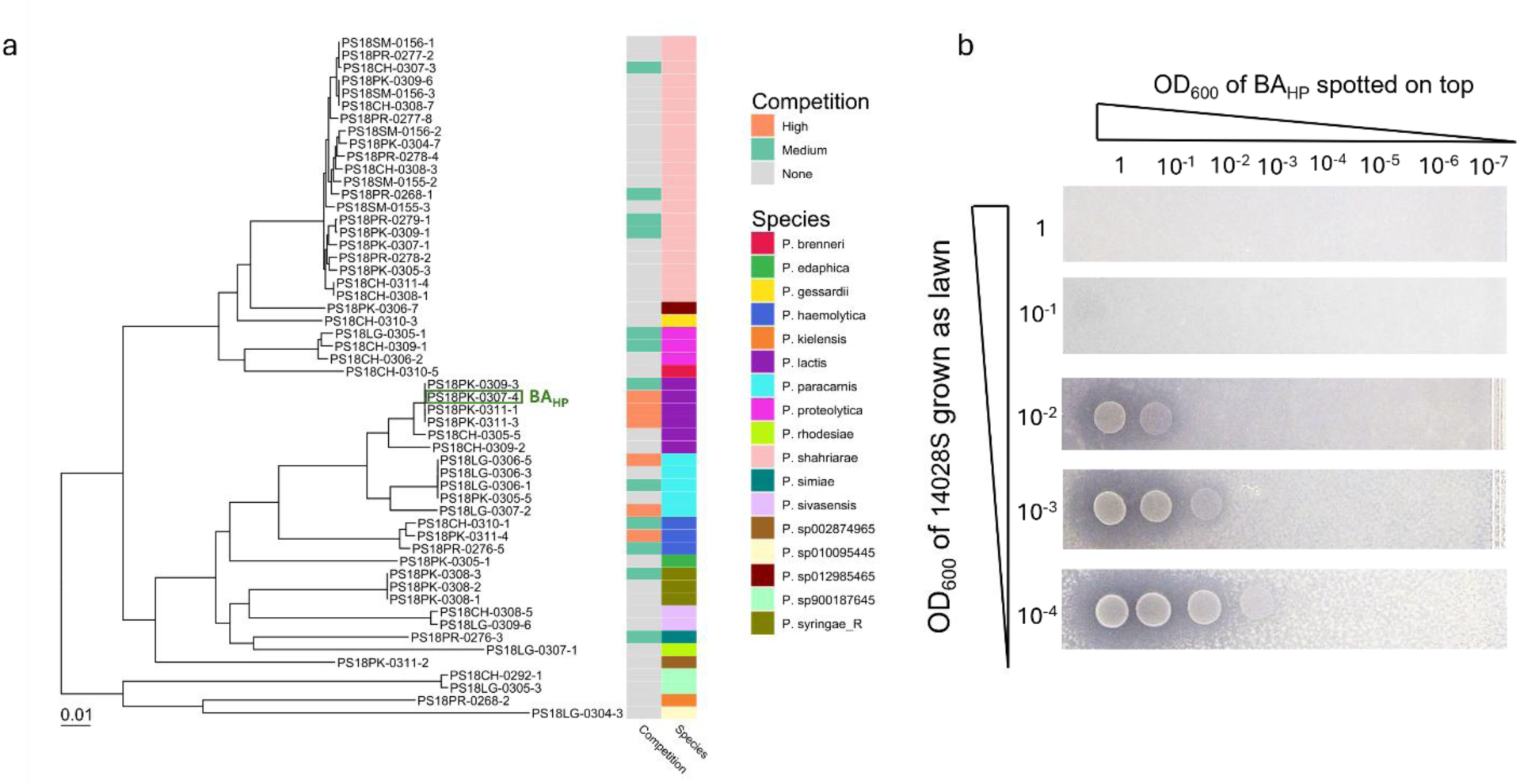
**a.** Maximum likelihood tree from core gene alignement of 54 P. fluorescens strains isolated from food products. The strains were characterised for their activity against Salmonella. Strains exhibiting “High” activity (clearing zones extending beyond the Pseudomonas spot) are highlighted in orange, the strains with “Medium” activity (clearing zones tightly surrounding the spot) in green and no activity in gray. Genome Taxonomy Database (GTDB) was used to predict the species of the P. fluorescens strains. **b.** Competition on agar against a Salmonella overlay after a 48-hour incubation, clearing zones indicate inhibition of the pathogen.

Activity of the BA_HP_ was first tested *in vitro* using an overlay assay on agar plates (Figure 1b). After 48 hours, distinct clearing zones were observed on the *Salmonella* lawn surrounding the BA_HP_ spotted on agar. The radius of the clearing zones was dilution dependent.

To investigate how widespread its activity is, we assessed BA_HP_ activity on agar overlay assays against eight *Salmonella* serovars in total and eight *Escherichia coli* isolates from food products. BA_HP_ was highly active, producing distinct inhibition zones on agar against all species tested (Figure S1).

### *Pseudomonas* effectively competes with *Salmonella in planta* in an application-dependent manner

To test whether or not BA_HP_ can compete with *Salmonella in planta*, we used a fresh produce alfalfa microgreen colonisation model we previously established (*19*). To get a better understanding of the dynamics of the interaction between BA_HP_ and *Salmonella in planta*, we tested their interaction using three inoculation strategies: co-inoculation, 48-hour pre-incubation with BA_HP_ and 48-hour pre-incubation with *Salmonella*. For the co-inoculation experiment, we normalised the inocula to achieve a 1:1 ratio (OD:600). We recovered and counted plant-associated bacterial cells every 24 hrs for three days using selective media for each strain to differentiate between BA_HP_ and *Salmonella*. We observed that when co-inoculated at a 1:1 ratio at the beginning, BA_HP_ recovery was consistently higher than *Salmonella* over the course of the experiment, with BA_HP_ becoming dominant in the population over time (Figure 2a). Under BA_HP_ pre-incubation, *Salmonella* numbers remained consistently low, while *Pseudomonas* followed an upwards growth trend (Figure 2b). When *Salmonella* was the first to be introduced to the plant system, BA_HP_ could still infiltrate and colonise the seedlings effectively, growing significantly and dominating the population over time, while *Salmonella* numbers remained stable (Figure 2c). In summary, in all application scenarios, BA_HP_ became the prevalent component of the population over time demonstrating a competitive advantage over *Salmonella*.

**Figure 2:**
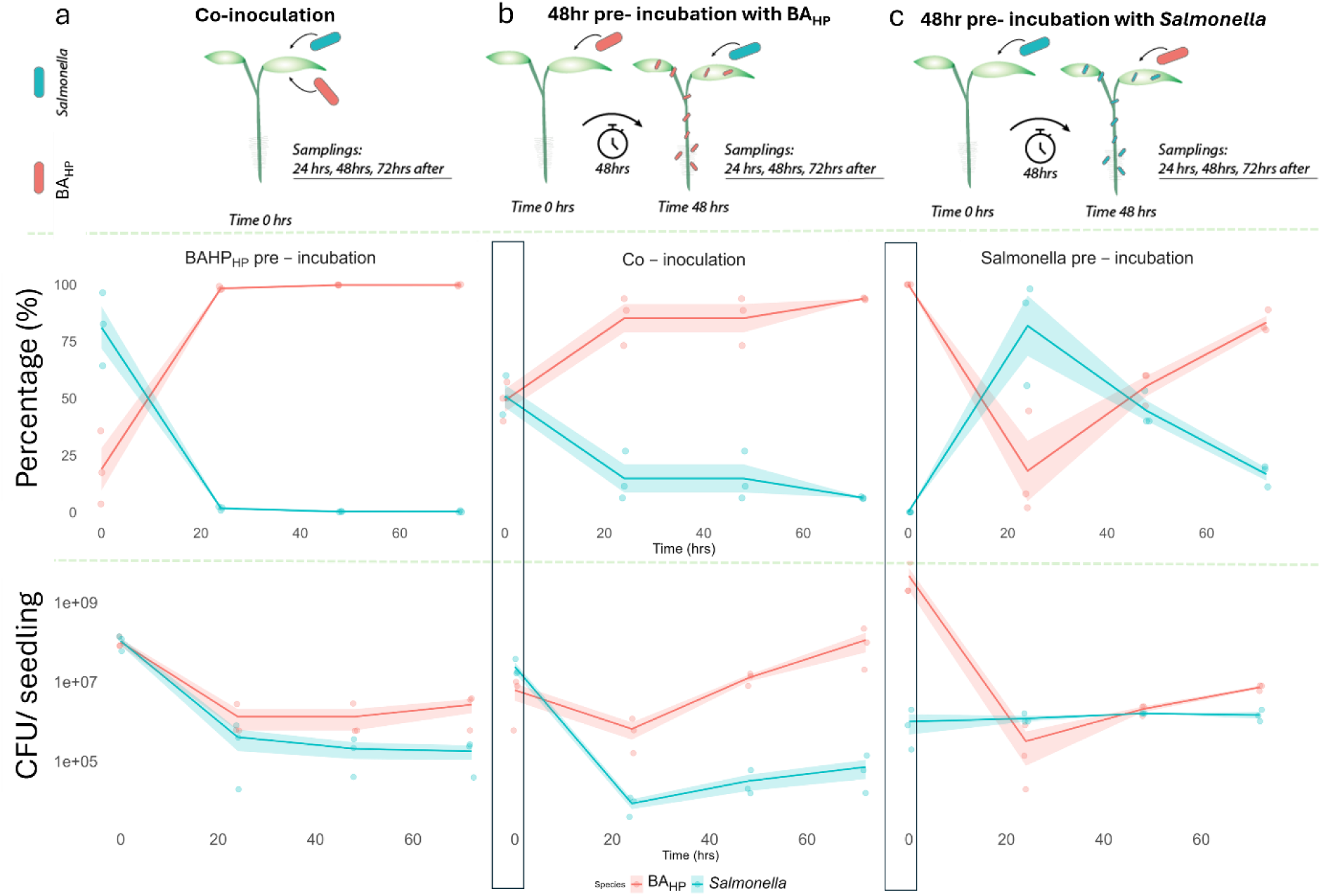
Illustration of relative abundance of Salmonella enterica Typhimurium 14028S and Pseudomonas fluorescens in planta over time following three inoculation strategies. Cells were recovered and counted every 24 hrs. Measurements are presented as a percentage (%) of each individual strain to the total population (graphs on top) and as cfu/ seedling (graphs on the bottom). **a.** BA_HP_ and Salmonella were inoculated at a 1:1 ratio and their growth was tracked over 72 hrs. BA_HP_ recovery emained consistently higher than Salmonella over the course of the experiment, establishing it as the dominant species within he population over time. **b.** BA_HP_ was introduced 48 hrs prior to Salmonella and recovery started 24 hrs after the introduction of Salmonella. Although Salmonella increased in numbers over time, BA_HP_ prior establishment and growth post Salmonella introduction established it as the dominant species within the population over time. **c.** Salmonella was introduced 48 hrs prior o BA_HP_ and cell recovery started 24 hrs after the introduction of BA_HP_. Although Salmonella was pre-introduced to the plant model, BA_HP_ increased rapidly in numbers, establishing as the dominant species within the population by 72hrs. BA_HP_ is shown n pink and Salmonella in blue. The lines represent the mean cfu.seedling / percentage of three biological replicates over time for each species. The shaded area around the lines represents the standard error of the mean. Dots represent the individual data points per timepoint. Boxes indicate the inoculation point of the second strain, and the cfu/percentage of recovered cells of the pre-established strain at the point of introduction.

### The gene expression profile of *Pseudomonas* in response to *Salmonella* is environment dependent

To identify the genes underpinning the competitive interaction between BA_HP_ and *Salmonella in vitro* and *in planta*, we performed RNA sequencing in both environments.

For the *in vitro* experiments, BA_HP_ spots were isolated 48 hours post incubation with *Salmonella* on an overlay agar inhibition assay, after confirming the presence of inhibitory clearing zones (Figure 3a). For the *in planta* bacterial extractions, we inoculated plants with an initial 1:1 ratio of BA_HP_ to *Salmonella*, as our previous findings demonstrated that both species coexist in the plant setting (Figure 2a). RNA was extracted 72 hrs post inoculation after confirming competition between the species by cell recovery and cfu counting (data not shown). To selectively isolate bacterial cells from plants, we optimised a gradient based isolation method using Gentodenz (*20*) (Figure 3b). This allowed the selective extraction of high-quality bacterial RNA from plant tissues. Three independent biological replicates were collected for each condition in each environment, leading to twelve samples in total.

**Figure 3:**
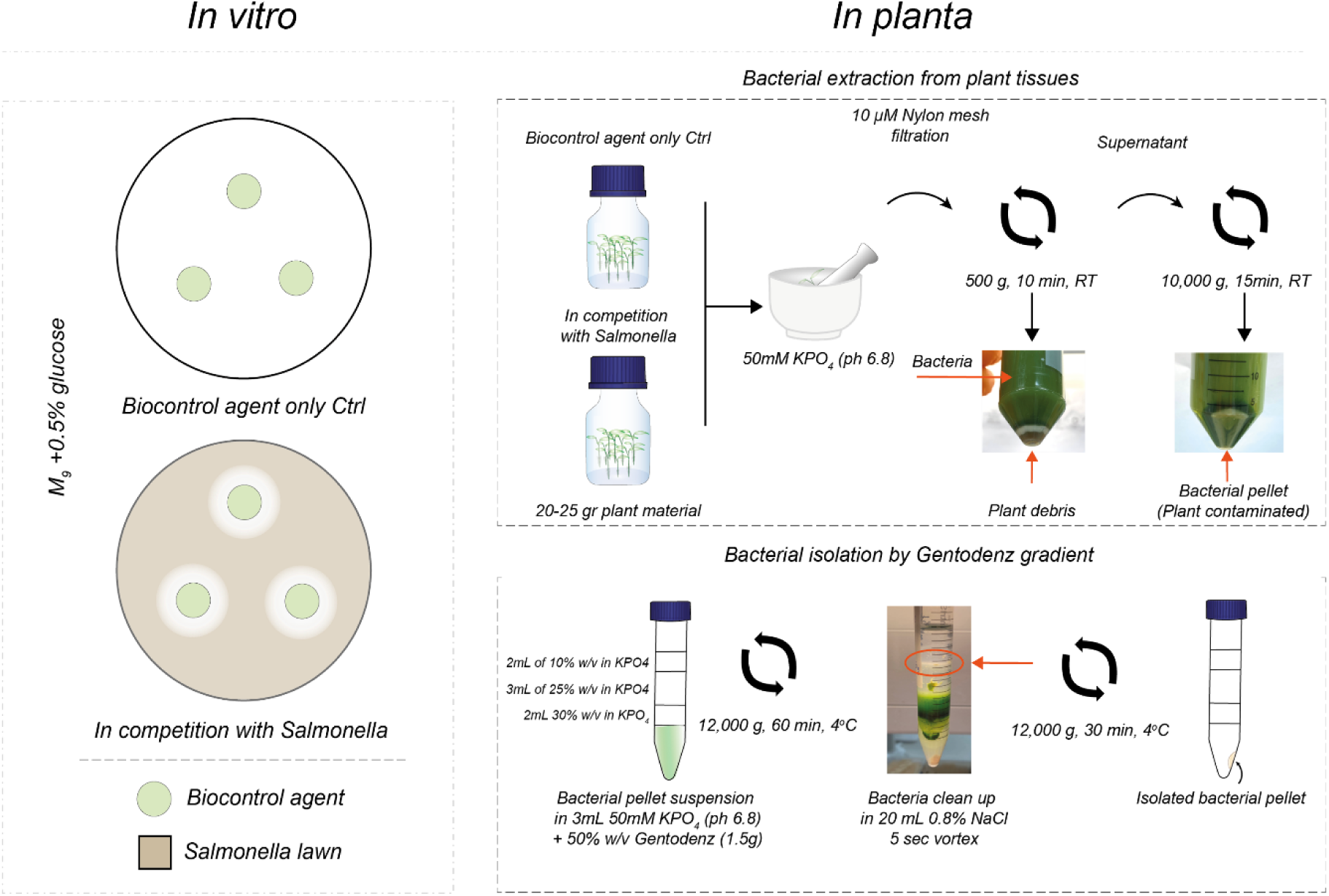
Schematic overview of the RNA extraction protocol performed in vitro and in planta.

Differential expression analysis was conducted by Deseq2, testing the impact of *Salmonella* on BA_HP_ in each environment (*in planta* vs *in vitro*). Each dataset was analysed for correlation with its corresponding replicates. The three *in planta* replicates were conducted on different days and analysis revealed that one replicate differed notably from the other two (Figure S3). To account for this variation, we analysed the data both with and without this replicate. Differentially expressed transcripts were considered statistically significant at a Benjamini-Hochberg corrected p value of <0.05 and an estimated log2 fold change (FC) ≥1 or ≤ −1 (Figure 4a-b, Tables 1-4). The effect of *Salmonella* presence in BA_HP_ differed markedly in the two environments. When tested *in vitro*, a large number of genes were identified as upregulated (N=237) or downregulated (N=416), with substantial fold-changes in expression and very consistent estimates of fold change across biological replicates (Figure 4a, Tables 1-2). In contrast, the *in planta* experiments (from the two replicate analysis) showed far fewer genes identified as significantly upregulated (N=27) or downregulated (N=59) (Figure 4b, Tables 3-4).

**Table 1:**

| Top 40 genes upregulated in vitro |  |  |  |  |  |
| --- | --- | --- | --- | --- | --- |
| Locus Tag | Name | Base Mean | Log2 Fold Change | p-value | Adj. p-value |
| P_04157 | dht | 7173.621 | 4.140 | 3.452e-287 | 9.904e-284 |
| P_01608 | bfr_1 | 8049.625 | 3.947 | 1.008e-251 | 1.928e-248 |
| P_04159 | preA | 2406.766 | 3.942 | 2.283e-210 | 2.620e-207 |
| P_03988 | araC | 10984.136 | 3.717 | 2.466e-104 | 3.824e-102 |
| P_03987 | ttuB_3 | 3199.325 | 3.638 | 4.192e-153 | 1.503e-150 |
| P_03985 | NA | 4194.789 | 3.621 | 8.393e-228 | 1.204e-224 |
| P_03986 | NA | 3244.606 | 3.551 | 4.228e-207 | 3.473e-204 |
| P_04158 | preT | 1979.512 | 3.278 | 2.825e-164 | 1.158e-161 |
| P_03984 | NA | 2060.681 | 3.039 | 4.947e-136 | 1.183e-133 |
| P_02836 | mdeA | 9978.615 | 2.910 | 2.630e-188 | 1.412e-185 |
| P_03983 | araF_1 | 1423.599 | 2.888 | 4.876e-101 | 6.825e-99 |
| P_03798 | antA | 659.028 | 2.698 | 8.016e-61 | 4.946e-59 |
| P_02043 | scoB | 2954.749 | 2.676 | 6.914e-92 | 7.629e-90 |
| P_02044 | scoA | 8097.997 | 2.577 | 4.054e-127 | 7.753e-125 |
| P_02042 | phaA | 4874.813 | 2.458 | 3.270e-64 | 2.108e-62 |
| P_03981 | rbsC_3 | 1266.000 | 2.424 | 1.630e-75 | 1.264e-73 |
| P_03979 | NA | 2612.361 | 2.364 | 2.464e-80 | 2.244e-78 |
| P_05649 | bauC_2 | 3028.744 | 2.310 | 3.636e-100 | 4.852e-98 |
| P_04356 | phhA | 5326.818 | 2.212 | 4.749e-75 | 3.585e-73 |
| P_02134 | NA | 23676.758 | 2.160 | 5.081e-81 | 4.703e-79 |
| P_03800 | antC | 190.587 | 2.159 | 6.666e-15 | 6.676e-14 |
| P_05303 | NA | 874.838 | 2.120 | 7.481e-37 | 2.247e-35 |
| P_02226 | fadJ | 2214.672 | 2.100 | 1.886e-112 | 3.279e-110 |
| P_04824 | catI | 752.097 | 2.077 | 3.680e-34 | 9.731e-33 |
| P_05648 | NA | 4166.786 | 2.069 | 4.251e-39 | 1.470e-37 |
| P_03980 | mro | 1830.928 | 1.998 | 6.084e-38 | 2.018e-36 |
| P_02225 | fadA_3 | 1927.641 | 1.984 | 9.773e-85 | 9.346e-83 |
| P_00219 | argD_1 | 1425.710 | 1.977 | 5.656e-69 | 3.958e-67 |
| P_04156 | pucI_2 | 2393.046 | 1.947 | 2.027e-51 | 9.774e-50 |
| P_04358 | aspC_2 | 9864.846 | 1.940 | 3.477e-76 | 2.771e-74 |
| P_03982 | araG_2 | 2901.814 | 1.929 | 5.493e-68 | 3.797e-66 |
| P_03793 | catA | 939.645 | 1.905 | 9.780e-30 | 2.218e-28 |
| P_04155 | hyuC_2 | 1887.495 | 1.882 | 9.163e-71 | 6.572e-69 |
| P_03802 | kynA | 1415.077 | 1.857 | 2.956e-48 | 1.357e-46 |
| P_04577 | NA | 2335.360 | 1.845 | 5.156e-59 | 3.082e-57 |
| P_04357 | NA | 819.673 | 1.841 | 5.451e-36 | 1.564e-34 |
| P_03790 | gabP_2 | 602.248 | 1.772 | 4.984e-37 | 1.537e-35 |
| P_03791 | kynU | 595.457 | 1.753 | 8.475e-37 | 2.520e-35 |
| P_01028 | NA | 2038.801 | 1.734 | 1.062e-36 | 3.125e-35 |

**Table 2:**

| Top 40 genes downregulated in vitro |  |  |  |  |  |
| --- | --- | --- | --- | --- | --- |
| Locus Tag | Name | Base Mean | Log2 Fold Change | p-value | Adj. p-value |
| P_04594 | mbtH | 1554.775 | -5.468 | 6.516e-203 | 4.673e-200 |
| P_05434 | pvdA | 5400.340 | -5.200 | 2.317e-193 | 1.477e-190 |
| P_03652 | NA | 1818.582 | -4.844 | 8.374e-136 | 1.922e-133 |
| P_00020 | trpC_1 | 1926.324 | -4.669 | 5.075e-130 | 1.004e-127 |
| P_03369 | NA | 1453.232 | -4.582 | 4.236e-207 | 3.473e-204 |
| P_04595 | dat | 2240.252 | -4.460 | 6.683e-20 | 9.331e-19 |
| P_03650 | pkn1 | 763.755 | -4.296 | 9.968e-134 | 2.200e-131 |
| P_01992 | hasA | 3780.430 | -3.932 | 9.499e-09 | 5.312e-08 |
| P_01660 | metE | 7021.946 | -3.866 | 3.671e-98 | 4.681e-96 |
| P_03835 | prsD_2 | 1421.656 | -3.860 | 3.021e-174 | 1.334e-171 |
| P_03645 | tycB_2 | 457.428 | -3.838 | 1.720e-54 | 9.137e-53 |
| P_05790 | NA | 1072.086 | -3.780 | 1.335e-78 | 1.161e-76 |
| P_03367 | lgrB | 13514.862 | -3.776 | 1.131e-28 | 2.430e-27 |
| P_02798 | ttgC | 1332.445 | -3.731 | 4.399e-141 | 1.147e-138 |
| P_00032 | trpD_1 | 956.826 | -3.718 | 3.227e-149 | 9.746e-147 |
| P_00803 | fumC_1 | 1026.018 | -3.659 | 4.822e-98 | 6.015e-96 |
| P_00028 | NA | 230.540 | -3.656 | 2.226e-52 | 1.101e-50 |
| P_03368 | lgrE | 1775.169 | -3.651 | 6.701e-104 | 1.012e-101 |
| P_04723 | znuB_2 | 393.411 | -3.634 | 1.948e-36 | 5.673e-35 |
| P_04725 | ssaB | 547.106 | -3.501 | 3.671e-83 | 3.453e-81 |
| P_00802 | NA | 477.735 | -3.499 | 7.941e-66 | 5.238e-64 |
| P_02799 | NA | 2568.008 | -3.498 | 5.452e-142 | 1.490e-139 |
| P_04724 | znuC_3 | 785.684 | -3.490 | 4.517e-65 | 2.945e-63 |
| P_00770 | NA | 209.028 | -3.462 | 2.443e-22 | 3.994e-21 |
| P_04499 | dltA_3 | 17284.653 | -3.411 | 1.237e-25 | 2.344e-24 |
| P_05789 | NA | 855.897 | -3.387 | 5.396e-78 | 4.553e-76 |
| P_00029 | trpB_1 | 3368.081 | -3.282 | 1.411e-183 | 6.746e-181 |
| P_04498 | NA | 1302.493 | -3.263 | 6.387e-50 | 2.979e-48 |
| P_04728 | NA | 369.626 | -3.251 | 3.396e-43 | 1.382e-41 |
| P_00025 | lysK | 1660.741 | -3.224 | 2.835e-149 | 9.037e-147 |
| P_02609 | NA | 131.043 | -3.223 | 1.360e-23 | 2.365e-22 |
| P_03833 | tolC_3 | 712.720 | -3.211 | 6.948e-97 | 8.306e-95 |
| P_00242 | fecI_1 | 552.146 | -3.173 | 7.836e-75 | 5.840e-73 |
| P_04729 | NA | 1179.416 | -3.160 | 1.449e-102 | 2.131e-100 |
| P_03646 | dltA_2 | 29246.572 | -3.149 | 3.184e-86 | 3.150e-84 |
| P_00027 | iolG_1 | 917.412 | -3.120 | 6.396e-101 | 8.739e-99 |
| P_00617 | efeU | 2023.128 | -3.114 | 1.831e-136 | 4.568e-134 |
| P_04634 | ugpA | 1449.929 | -3.093 | 1.717e-150 | 5.795e-148 |
| P_01607 | NA | 2009.016 | -3.071 | 2.127e-132 | 4.359e-130 |
| P_04727 | NA | 260.638 | -3.039 | 5.307e-35 | 1.457e-33 |

**Table 3:**

| Top 40 genes upregulated in planta |  |  |  |  |  |
| --- | --- | --- | --- | --- | --- |
| Locus Tag | Name | Base Mean | Log2 Fold Change | p-value | Adj. p-value |
| P_03958 | NA | 1185.514 | 1.595 | 2.146e-04 | 2.355e-03 |
| P_03449 | NA | 572.993 | 1.344 | 3.593e-09 | 2.165e-07 |
| P_03739 | NA | 2915.672 | 1.335 | 1.142e-08 | 5.979e-07 |
| P_03450 | NA | 3301.042 | 1.215 | 3.513e-16 | 8.667e-14 |
| P_04747 | NA | 5966.631 | 1.200 | 1.461e-22 | 9.462e-20 |
| P_03960 | nemA_2 | 30799.651 | 1.190 | 1.143e-07 | 4.626e-06 |
| P_01878 | ompW | 9104.110 | 1.174 | 4.038e-19 | 1.743e-16 |
| P_05120 | arcD_1 | 14535.527 | 1.171 | 1.219e-19 | 6.317e-17 |
| P_03451 | ccoN1_2 | 7118.683 | 1.159 | 3.699e-19 | 1.742e-16 |
| P_04185 | deaD | 30130.348 | 1.122 | 3.040e-08 | 1.389e-06 |
| P_03441 | NA | 13461.587 | 1.105 | 1.384e-27 | 1.434e-24 |
| P_03558 | adhB_2 | 5658.160 | 1.098 | 6.656e-17 | 1.916e-14 |
| P_00861 | nemR_2 | 8558.231 | 1.091 | 1.477e-11 | 1.780e-09 |
| P_03623 | NA | 483.455 | 1.087 | 4.393e-07 | 1.450e-05 |
| P_03448 | ccoP1 | 5663.769 | 1.051 | 3.337e-17 | 1.081e-14 |
| P_00860 | yqhD | 30171.838 | 1.050 | 8.429e-04 | 7.218e-03 |
| P_01240 | NA | 1650.526 | 0.990 | 3.515e-11 | 3.875e-09 |
| P_03440 | NA | 2685.222 | 0.989 | 1.031e-14 | 2.054e-12 |
| P_02897 | NA | 1031.892 | 0.973 | 7.228e-07 | 2.177e-05 |
| P_02898 | roxA | 5089.504 | 0.972 | 1.461e-12 | 2.046e-10 |
| P_01814 | uctC_2 | 6631.014 | 0.956 | 2.394e-09 | 1.531e-07 |
| P_01813 | mmgC_2 | 5672.012 | 0.955 | 1.376e-08 | 7.060e-07 |
| P_03962 | nemR_3 | 22022.739 | 0.949 | 2.139e-03 | 1.552e-02 |
| P_00859 | hchA_1 | 31402.699 | 0.948 | 2.021e-05 | 3.537e-04 |
| P_05530 | NA | 4691.642 | 0.946 | 2.707e-05 | 4.424e-04 |
| P_05532 | NA | 782.649 | 0.929 | 2.854e-11 | 3.286e-09 |
| P_05397 | spuE_2 | 728.872 | 0.910 | 1.601e-04 | 1.869e-03 |
| P_01531 | NA | 626.230 | 0.908 | 1.757e-07 | 6.646e-06 |
| P_04195 | curA_2 | 18419.600 | 0.902 | 2.109e-03 | 1.535e-02 |
| P_05763 | rimO_2 | 11774.705 | 0.882 | 8.675e-09 | 4.633e-07 |
| P_05531 | rhaS_12 | 5512.839 | 0.879 | 1.289e-06 | 3.425e-05 |
| P_04628 | argO_2 | 32570.162 | 0.875 | 1.045e-09 | 7.317e-08 |
| P_05301 | sotB_1 | 1287.895 | 0.864 | 1.662e-09 | 1.090e-07 |
| P_05289 | gbuA | 1986.675 | 0.862 | 3.740e-05 | 5.890e-04 |
| P_04807 | plsX | 3067.950 | 0.854 | 1.088e-09 | 7.513e-08 |
| P_03439 | copA_1 | 4074.284 | 0.852 | 5.638e-09 | 3.210e-07 |
| P_01357 | thil | 8113.961 | 0.848 | 3.001e-10 | 2.508e-08 |
| P_00475 | betT2 | 8242.898 | 0.838 | 6.257e-13 | 9.535e-11 |
| P_04921 | ccmB | 643.550 | 0.837 | 6.503e-08 | 2.808e-06 |
| P_05575 | rhaS_13 | 5890.236 | 0.836 | 1.587e-10 | 1.394e-08 |

**Table 4:**

| Top 40 genes downregulated in planta |  |  |  |  |  |
| --- | --- | --- | --- | --- | --- |
| Locus Tag | Name | Base Mean | Log2 Fold Change | p-value | Adj. p-value |
| P_04371 | dksA_2 | 101.712 | -2.954 | 3.882e-24 | 3.352e-21 |
| P_04370 | yciC_2 | 750.311 | -2.284 | 4.885e-28 | 6.327e-25 |
| P_02142 | copZ | 1201.361 | -1.700 | 5.010e-08 | 2.200e-06 |
| P_01673 | NA | 2391.785 | -1.664 | 3.250e-32 | 8.419e-29 |
| P_03988 | araC | 1373.425 | -1.660 | 2.000e-15 | 4.317e-13 |
| P_02959 | mgtB | 19101.142 | -1.601 | 7.729e-14 | 1.292e-11 |
| P_04369 | NA | 226.893 | -1.540 | 1.044e-15 | 2.351e-13 |
| P_01674 | NA | 3393.599 | -1.540 | 4.787e-41 | 2.480e-37 |
| P_04373 | amiC_3 | 773.105 | -1.493 | 2.159e-23 | 1.598e-20 |
| P_01882 | NA | 1061.167 | -1.419 | 2.311e-09 | 1.497e-07 |
| P_04516 | NA | 9702.047 | -1.418 | 5.600e-32 | 9.670e-29 |
| P_04645 | NA | 17573.363 | -1.355 | 1.751e-16 | 4.629e-14 |
| P_02139 | hmrR | 1192.167 | -1.343 | 3.579e-06 | 7.923e-05 |
| P_04482 | hcaB_7 | 1989.930 | -1.332 | 9.915e-19 | 3.951e-16 |
| P_02065 | mgtC | 3166.099 | -1.330 | 1.324e-08 | 6.859e-07 |
| P_05584 | tcyC | 820.210 | -1.326 | 1.322e-10 | 1.246e-08 |
| P_02960 | NA | 957.246 | -1.310 | 1.211e-05 | 2.274e-04 |
| P_01869 | NA | 6280.976 | -1.276 | 2.320e-18 | 8.015e-16 |
| P_01740 | fgd_2 | 52231.175 | -1.268 | 9.192e-22 | 5.292e-19 |
| P_01870 | NA | 1858.746 | -1.263 | 6.777e-13 | 1.003e-10 |
| P_01084 | ibpA | 17884.591 | -1.255 | 4.186e-17 | 1.276e-14 |
| P_00572 | dxs_1 | 4095.617 | -1.248 | 1.532e-04 | 1.813e-03 |
| P_03987 | ttuB_3 | 1031.568 | -1.232 | 1.893e-13 | 3.065e-11 |
| P_01692 | NA | 1052.451 | -1.213 | 3.169e-06 | 7.401e-05 |
| P_01675 | NA | 821.188 | -1.180 | 1.688e-10 | 1.457e-08 |
| P_04385 | NA | 232.036 | -1.180 | 8.179e-07 | 2.381e-05 |
| P_01397 | NA | 16267.228 | -1.171 | 1.120e-12 | 1.612e-10 |
| P_04075 | clpB | 230550.544 | -1.142 | 7.535e-10 | 5.741e-08 |
| P_00571 | NA | 635.157 | -1.139 | 7.071e-08 | 3.028e-06 |
| P_00656 | osmY_2 | 111809.018 | -1.099 | 1.153e-18 | 4.267e-16 |
| P_01083 | NA | 529.534 | -1.084 | 3.113e-06 | 7.332e-05 |
| P_03169 | fabG_4 | 3823.295 | -1.070 | 3.923e-12 | 5.212e-10 |
| P_00951 | NA | 358.024 | -1.063 | 1.170e-04 | 1.454e-03 |
| P_04431 | ltaE_2 | 6600.561 | -1.061 | 5.221e-16 | 1.229e-13 |
| P_04774 | ohrB_2 | 14503.910 | -1.055 | 1.101e-03 | 9.160e-03 |
| P_01660 | metE | 967.062 | -1.048 | 6.456e-07 | 2.003e-05 |
| P_02961 | NA | 1560.957 | -1.047 | 5.024e-05 | 7.501e-04 |
| P_03656 | NA | 9152.936 | -1.037 | 1.167e-09 | 7.937e-08 |
| P_00339 | argD_2 | 1783.680 | -1.035 | 8.129e-05 | 1.129e-03 |
| P_03125 | leuD | 15004.037 | -1.035 | 2.973e-06 | 7.066e-05 |

**Figure 4:**
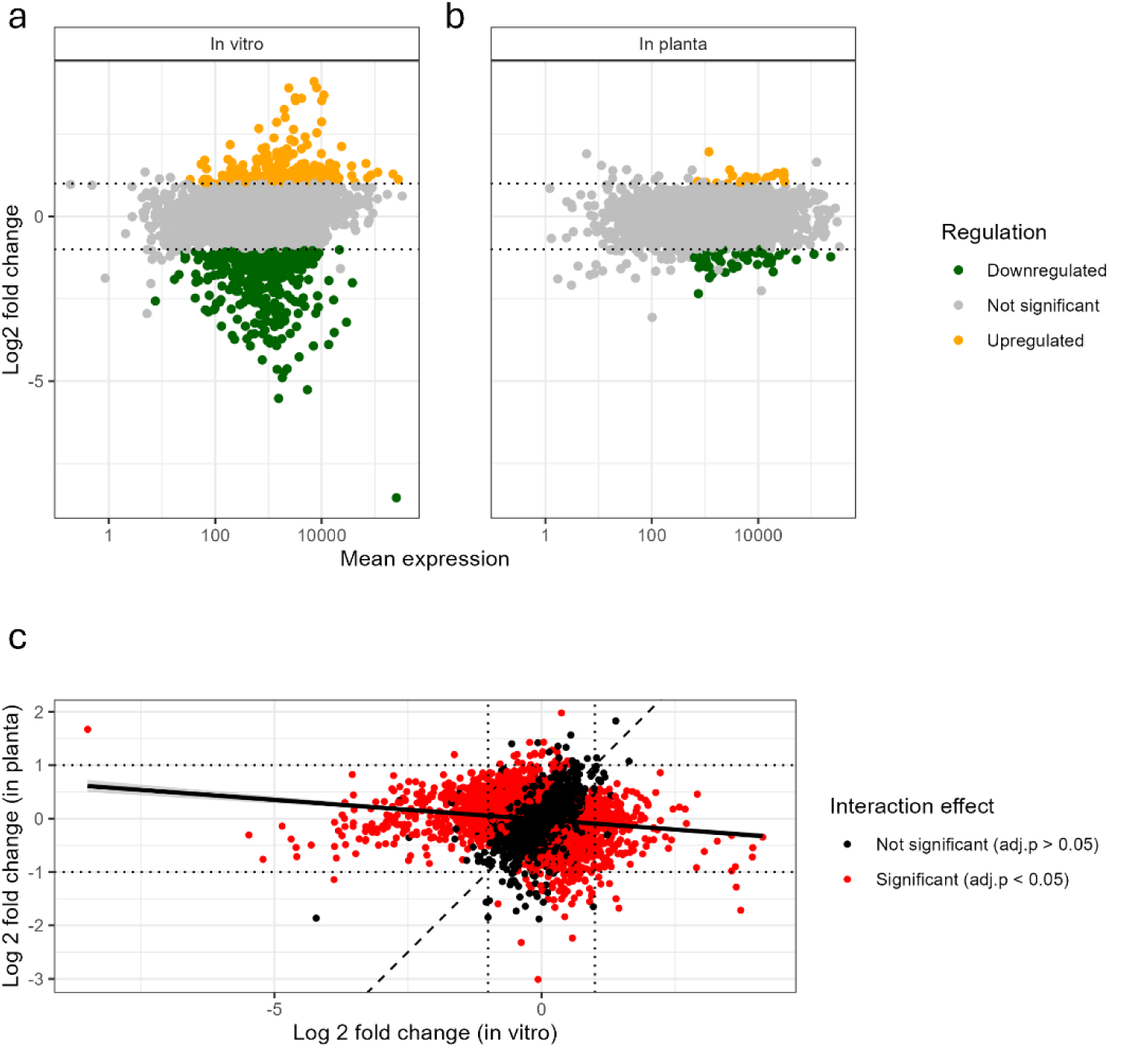
RNA sequencing results summary. **a-b**: MA plots (log2 fold change vs. mean expression) showing the gene expression changes in response to Salmonella in vitro (a) and in planta (b). Changes in expression was only considered significant when adj.p<0.05 and LogFC ≥ 1/≤ −1. Green dots represent downregulated genes, while orange dots represent upregulated genes. Grey dots are not significant. **c.** Scatter plot showing the relationship between log2 fold changes in gene expression under in vitro and in planta conditions. Each point represents a gene, with its log2 fold change in vitro on the x-axis and in planta on the y-axis. The solid black line indicates the best-fit linear regression, showing the overall trend. The dotted black line represents the identity line (y = x), where genes would fall if their expression changed equally in both conditions. Red points highlight genes with a statistically significant deviation (adj.p<0.05) from the expected relationship.

To test the effect of the environment on the transcriptional response of *Pseudomonas* to *Salmonella*, we compared the fold change of each gene in each biological pair in both conditions (presence and absence of *Salmonella*) and in both environments (*in vitro* and *in planta*). We found there was a small negative correlation (Spearman’s rho= −0.2, p<0.001) between the fold-changes estimated *in vitro* vs those estimated *in planta* (Fig 4c), and a slight negative concordance between the genes identified as upregulated vs downregulated in each environment (Kendall’s tau-b = −0.03, 95% CI= −0.06 to −.003). There were a large number of genes (N=1084) with significant interaction effects suggesting a significantly different impact of *Salmonella* across environments. Ten genes were differentially expressed in both environments, but notably eight of these were in different directions (see Table 7).

Sensitivity analyses including all three *in vitro* and *in planta* replicate pairs confirmed the central observation; expression patterns were highly consistent and reproducible among *in vitro* replicates, and distinct from the *in planta* environment (Figure S2). Fewer significantly differentially expressed genes were identified *in planta* compared to *in vitro*. These results are consistent with the original findings, demonstrating that the BA_HP_ response to *Salmonella* is conditional, underscoring the inherent complexity and variability of the *in planta* experimental system.

### Key metabolic pathways in BA_HP_ are differentially enriched in the two environments in the presence of *Salmonella enterica* Typhimurium 14028S

To identify significant co-ordinated effects across gene sets grouped by their involvement in related biological functions or pathways, we performed pathway analysis using the KEGG pathway Ontology database (*21*).

Exposure to *Salmonella* resulted in the induction or repression of numerous genes within specific pathways in an environment-specific manner (Tables 5-6). Pathways related to flagellum biosynthesis, tryptophan metabolism and benzoate degradation were important in the *in vitro* environment, while in the *in planta* environment most changes were observed in central metabolism (Figure 5a, Tables 5-6). Notably, six pathways: degradation of aromatic compounds, porphyrin metabolism, oxidative phosphorylation, bacterial secretion SecA system, peptidoglycan synthesis, and tyrosine metabolism, were all enriched in both environments but the genes in these pathways showed different expression patterns. For example, genes within the peptidoglycan synthesis pathway were downregulated *in vitro* but upregulated *in planta* (Figure 5b). Overall, this transcriptomic analysis highlights each environment affects *Pseudomonas* gene expression differently in the presence of *Salmonella enterica* Typhimurium 14028S.

**Table 5:**
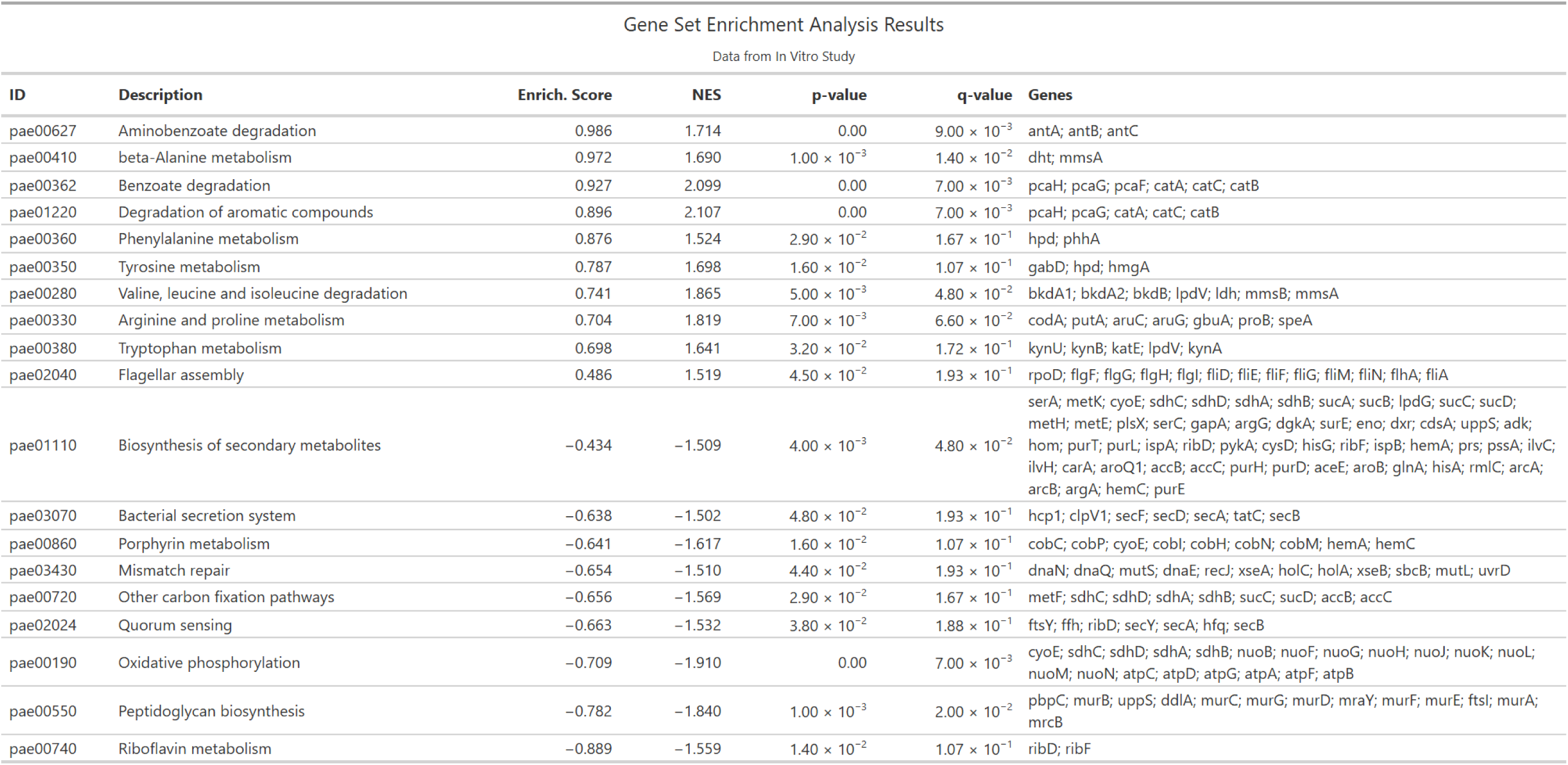

**Table 6:**
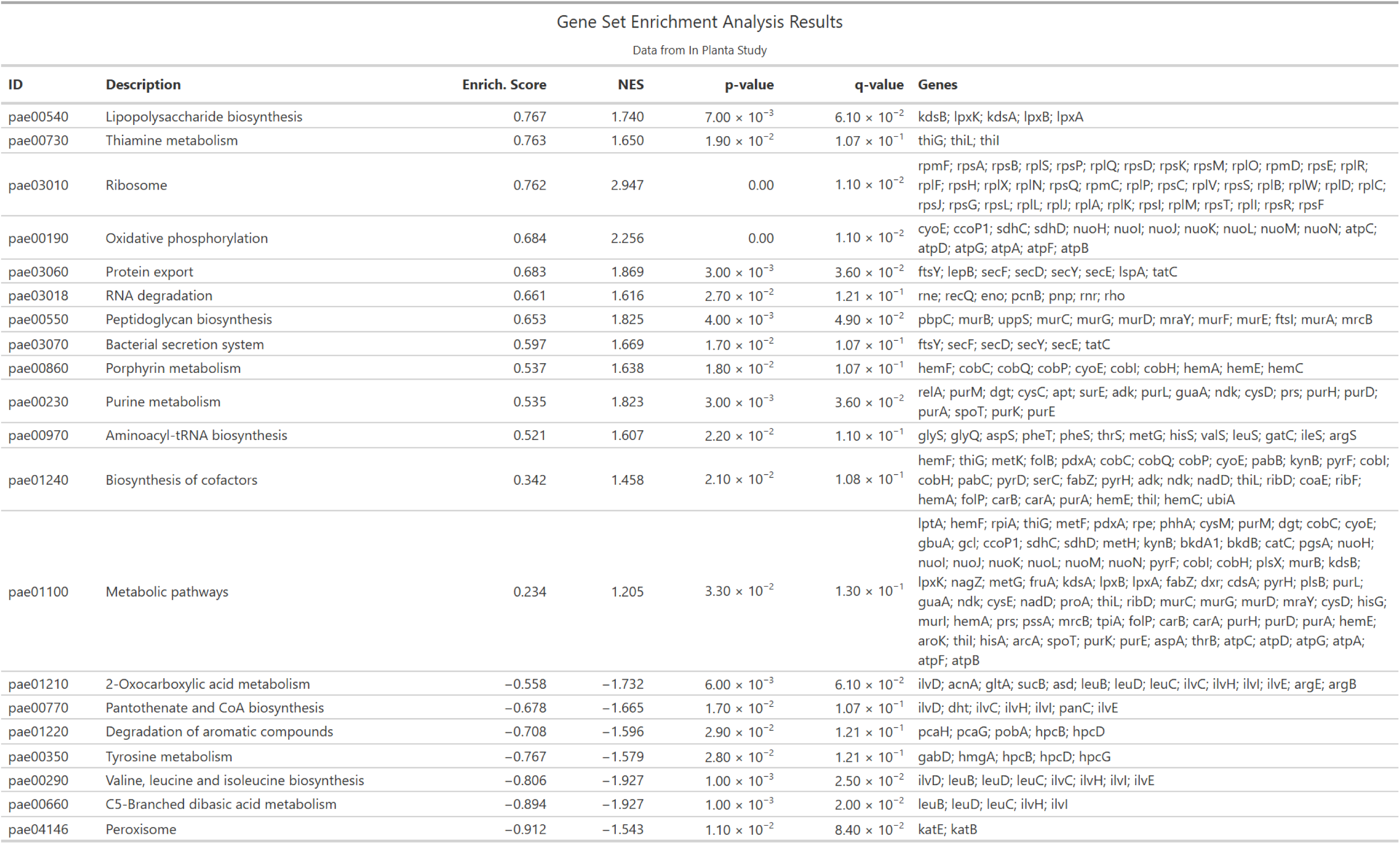

**Table 7:**

| Overlap of Differential Expression |  |  |  |
| --- | --- | --- | --- |
| In Vitro vs In Planta |  |  |  |
| Gene Name | Locus Tag | In Vitro | In Planta |
| NA | P_00571 | Up | Down |
| dxs_1 | P_00572 | Up | Down |
| nemR_2 | P_00861 | Up | Up |
| NA | P_01397 | Up | Down |
| metE | P_01660 | Down | Down |
| NA | P_01673 | Up | Down |
| NA | P_03656 | Up | Down |
| ttuB_3 | P_03987 | Up | Down |
| araC | P_03988 | Up | Down |
| arcD_1 | P_05120 | Down | Up |

**Figure 5:**
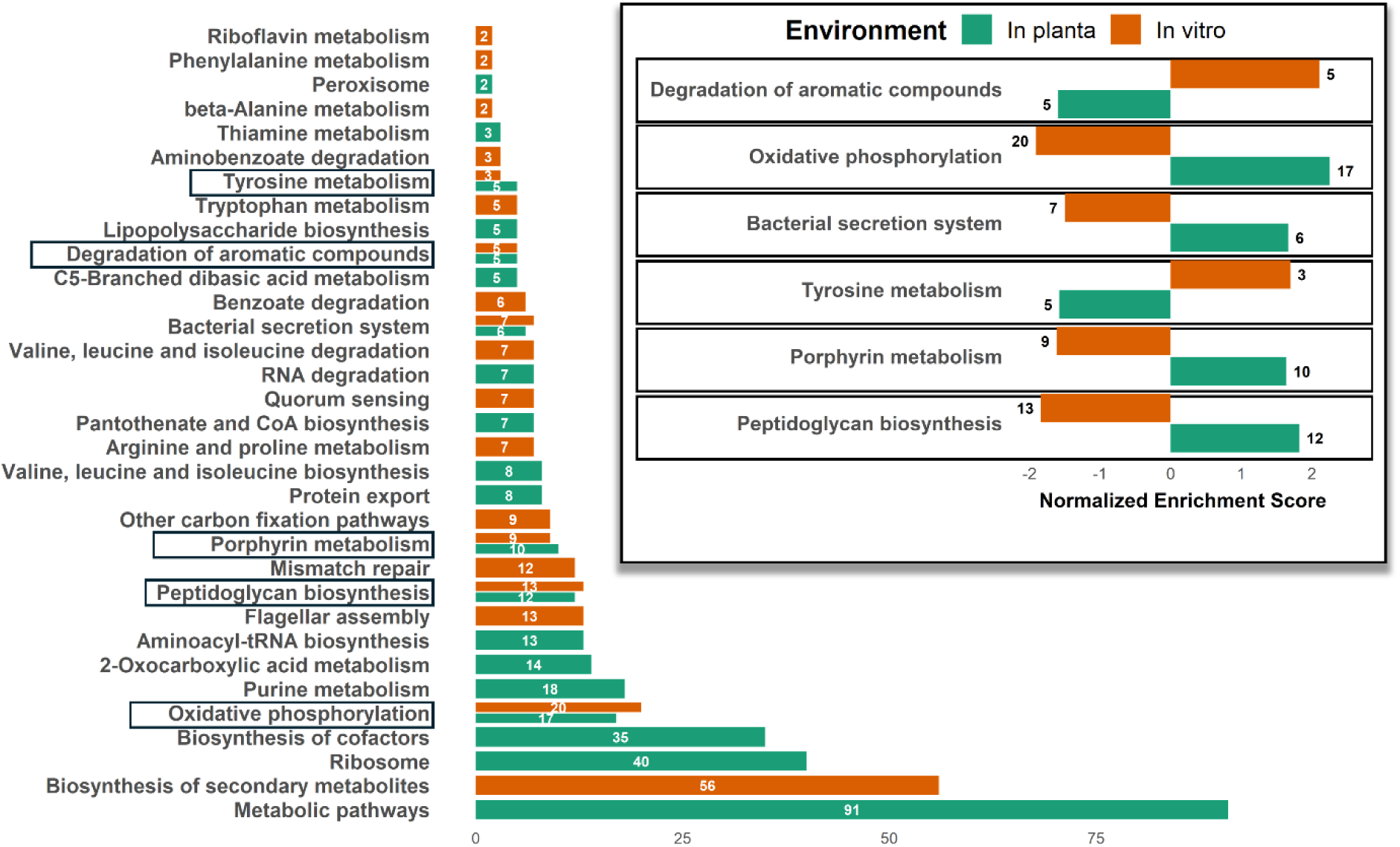
**a.** Pathway enrichment analysis based on the KEGG pathway Ontology database on BA_HP_ in the presence of Salmonella enterica Typhimurium. The X axis shows the number of core genes that contribute the most to enrichment score per pathway, and the Y axis lists the significantly enriched pathways (by KEGG). The in planta environment is represented in green and the in vitro in orange. The numbers in white represent the number of differentially expressed genes within each pathway. Highlighted in black boxes are pathways shared between environments**. Inset: b.** Shared pathways between environments. The X axis shows the normalised enrichment score. Directionality indicates positive or negative expression of the genes within the pathways with Up being indicated as +ve values on the x axis or down indicated in -ve values on the X axis.

## Materials and Methods

### Strains and growth media

Strains used in this study were *Salmonella enterica subsp. enterica* serovar Typhimurium strain 14028S tagged with a kanamycin resistance cassette and the lacZ operon (14028S::lacIZ) (*22*) to facilitate selection in competition experiments. The *Salmonella* strains used to demonstrate the wide range of activity of BA_HP_ (supplementary Figure S1) were kindly gifted by Professor Rob Kingsley (Quadram Institute). The *Escherichia coli* isolates used in the same assay were originally isolated from food products by the Mather group (*23*) (Quadram Institute). The selected *P. fluorescens* strain (BA_HP_) used in subsequent competition experiments and RNA sequencing belongs to the *Pseudomonas lactis* species (PS18PK-0307-4 in bio project PRJNA973713). *Salmonella* was differentiated by *Pseudomonas* by growth on selective media (LB agar supplemented with 40 μg/ mL Xgal, 1mM IPTG, and 50 μg/ mL kanamycin for *Salmonella* or Cetrimide, Fucidin, and Cephaloridine (CFC) *Pseudomonas* selective agar for *Pseudomonas*).

### Phylogenetic analysis

The 54 *P. fluorescens* isolates were previously cultured from retail food from Norfolk and whole genome sequenced (*18*). Whole genome sequencing data for the strains is uploaded to the Sequence Read Archive under bio projects PRJNA973713 and PRJNA1248571. The sequence for the selected *Pseudomonas* BA_HP_ can be found under the name PS18PK-0307-4. The Illumina were trimmed using Fastp v0.19.5 (*24*), assembled using Spades v1.1.0 (*25*) and the species was predicted using GTDB v2.2.2 (*26*). Assemblies were annotated using BAKTA v1.9.3 (*27*) and clustered using Roary v3.10.2 (*28*) with 95% identity cut-offs. Genes were regarded as core if they were found in 95% of isolates and RAxML v8.2.4 (*29*) was used to form a maximum-likelihood tree from the core gene alignment and GTDB was used to predict the *P. fluorescens* species.

### Competition assays

For the *in vitro* competition assay on agar, overnight cultures of *Pseudomonas* were set up in M9 media supplemented with 0.5% glucose or LB for *Salmonella* (14028S::lacIZ) supplemented with 50 μg/ mL kanamycin. *Salmonella* and *Pseudomonas* cultures were diluted to an optical density (OD) of 1 at 600 nm in PBS and then serially diluted to the required dilutions. 10 mL of appropriate *Salmonella* dilutions were poured on M9 agar plates (supplemented with 0.1% glucose) and left to settle on the surface for four minutes before discarding the excess liquid and allowed to dry for 30 minutes in a safety cabinet. 5 μL of appropriate *Pseudomonas* dilutions were spotted on the *Salmonella* lawn plates and were incubated at 20°C for 2 days. All procedures were conducted under sterile conditions in a class II microbiological safety cabinet.

For the *in planta* competition assays, alfalfa seeds were sterilised by immersion in 10 mL of 70% ethanol for 30 seconds, followed by three washes with sterile water and one wash with 10 mL of 5% sodium hypochlorite for 3 minutes. After five additional sterile water washes, seeds were transferred onto Murashige and Skoog plates to germinate and incubated in darkness at room temperature for 48 hours. Bacterial strains were grown overnight in appropriate growth media (*Pseudomonas* in M9 with 0.5% glucose or LB and *Salmonella* in LB) at 30°C the night before the inoculation (48 hours post sterilisation). For inoculating the seedlings, bacterial strains were diluted to OD 0.02 in 10mM MgCl_2_ and germinated seedlings of similar sizes were transferred to water agar. For the co-inoculation experiment the strains were premixed 1:1. A 5 μL inoculum was applied at the junction between the root and hypocotyl of each seedling, which were then incubated under plant growth lights (16-hour on/8-hour off cycle, simulating natural sunlight patterns), on water agar plates with different strains/strain combinations on separate plates. For the preincubation experiments, the second strain was introduced 48 hours later as a 5 μl spot of normalised O/N cultures to an OD of 0.02. Samplings were then carried out 24, 48, 72 hours after the second inoculation. For bacterial recovery from plant roots, seedlings were gently washed in PBS to remove any non-specifically attached cells, ground in autoclaved mortars with PBS, and serial dilutions were plated onto selective for each species agar plates. Plates were incubated at 30°C and cell counting was carried out 48 hrs later. Three biological replicates were included per inoculation scenario.

### RNA extractions and RNAseq analysis

For the agar competition experiments, RNA was extracted from *Pseudomonas* spots in competition with *Salmonella* on agar at 48 hours post plate inoculation. Three biological replicates were processed in two distinct conditions: presence and absence of *Salmonella* in the form of a lawn on the plates. Inhibition on the plates with *Salmonella* was confirmed by visual inspection before cells were harvested from the *Pseudomonas* spots.

Bacterial RNA extractions from inoculated plants were based on (*20*, *30*). Briefly, seeds were sterilised and germinated as described before and grown in 10 Duran bottles per replicate. 48 hours post germination they were co-inoculated with 1:1 mix of *Pseudomonas: Salmonella* normalised to an OD_600nm_ 0.2. After 72 hrs samples were taken to confirm competition and colonised plant tissue was processed as follows. 10-25 grams of plant tissue were disrupted in 50 mM potassium phosphate buffer in a sterile mortar and pestle (KPO_4_, pH 6.8). The disrupted tissue was incubated on a rotating wheel at 4°C for 1 hour in 50 mL Falcon tubes to a final KPO_4_ volume of 40 mLs. The resulting suspension was filtered through a sterile100 μM nylon mesh to remove coarse plant debris. The filtrate was centrifuged at 500 × *g* for 10 minutes at room temperature to pellet the remaining plant debris, retaining bacteria in the supernatant. The supernatant was collected and centrifuged at 10,000 × *g* for 15 minutes at room temperature to pellet bacteria along with any residual plant material. The pellet was resuspended in 3 mL of KPO_4_ buffer containing 50% w/v Gentodenz (1.5 g). A gel gradient was created by carefully layering 2 mL of 10% w/v Gentodenz in KPO_4_, 3 mL of 25% w/v Gentodenz in KPO4, and 2 mL of 30% w/v Gentodenz in KPO4 over the resuspended sample. The gradient was centrifuged at 12,000 × *g* for 60 minutes at 4°C. The bacterial layer (3–4 mL), as shown on Figure 3, was cleaned by adding 20 mL of 0.8% NaCl and vortexing briefly (5 seconds), and centrifuging again at 12,000 × *g* for 30 minutes at 4°C. Finally, the bacterial pellet was washed with RNA-free water and transferred to an Eppendorf tube for RNA extraction. RNA extractions were carried out using the RNeasy Plus Kits (Qiagen), followed by an extra DNAase treatment (Ambion TURBO).

RNA extractions were performed in triplicate for each condition. RNA was quantified using the Qubit RNA high Sensitivity kit (Thermo Fisher Scientific) and the quality was assessed using the High sensitivity RNA ScreenTapes on a Tapestation (Agilent). RNA sequencing was performed by Azenta Life Sciences, including library preparation, sequencing on an Illumina NovaSeq 6000 platform using a 2 × 150 bp paired-end configuration (Version 2.0.0), and initial quality control.

RNA-seq data were processed as paired-end reads. Analysis was performed as outlined in (*31*) using the Galaxy platform (*32*). Briefly, read quality was assessed using fastQC (*33*), followed by adapter trimming with fastp (*24*). Trimmed-paired end reads were aligned to the sequenced reference genome using HISAT2 (*34*), and the number of reads successfully mapped were quantified using samtools stats (*35*). Featurecounts (*36*) was used to assign mapped reads to annotated regions, and differential gene expression quantified using DESeq2 (*37*). Normalised counts estimated fold changes were compared for all biological replicates for an indication of consistency between pairs of samples. DESeq2 analysis was then conducted with the environment (*in vitro* vs *in planta*) as the main effect, the interaction between *Salmonella* competition and environment, and a nested ID variable corresponding to sample pair included in the model. The effects of *Salmonella* in each environment, and the test for the difference in fold chage between the environments (the interaction effect) were derived from this model. A sensitivity analysis was conducted including or excluding the first *in planta* replicate, and a further sensitivity analysis was conducted with DESeq2models estimated independently in each environment. Log2 Fold Changes were shrunk using the apeglm method (*38*). Dispersion plots, size factors and mean/SD plots were used as model diagnostics. Genes were considered differentially expressed if the Benjamini-Hochberg adjusted P-value reported by Deseq2 was below 0.05 and the Log2 fold change was ≥ 1 or ≤ −1. Graphs were generated using R using ggplot2 (*39*). Interaction effects were considered statistically significant when the adjusted p-value was less than 0.05. Full counts, code and output are supplied.

### Pathway analysis

For the pathway enrichment analysis we used the R Package ClusterProfiler (*40*) on the output of the differential expression analysis (performed by DESeq2) between controls and single conditions. A ranked gene list based on differential expression values was used as input, including all three *in vitro* replicates and the two more correlated *in planta*. This analysis used the KEGG Pathway Ontology database (*21*), and gene symbols were mapped to the *Pseudomonas* aeruginosa PAO1 reference genome, using the ‘pae’ organism annotation package within KEGG. PAO1 was selected due to it being the closest annotated reference with well-characterised pathways. The genes were ranked according to logFC and the ‘gseKegg’ function, within ClusterProfiler, to conduct a leading-edge analysis. The analysis was conducted with the following parameters: organism = “pae”, nPerm = 10000, minGSSize = 3, maxGSSize = 800, pvalueCutoff = 0.05, and pAdjustMethod = “none”. The KEGG gene identifiers were provided using the keyType = “ncbi-geneid”. The results consisted of an enrichment score (ES), a normalised enrichment score (NES) and the core genes that contribute the most to this ES per pathway. Directionality is given by a positive or negative ES indicating that genes were up or down regulated respectively across the pathway. The ES lies between −1 and 1. The NES is calculated by dividing the ES by the expected value of the ES score for a random gene set of the same size. The NES allows to compare the degree of enrichment of different gene sets to each other. We only considered pathways as significantly enriched that had a *p-*value of <0.05.

## Data Availability Statement

The raw data (full counts), and analysis code, that support the findings of this study are openly available in the GitHub repository at https://github.com/Eleftheria-trm/*Pseudomonas*_*Salmonella*_competition_2025. All files and scripts required to reproduce the results are provided.

## Acknowledgements

We thank Professor Rob Kingsley’s group (Quadram Institute) for providing the additional *Salmonella* serovars used to test the range of BA_HP_ activity in this study.

## Discussion

Several species within the *Pseudomonas* genus have been described to exhibit strong activity not only against plant pathogens (*8*, *41–43*) but also human pathogens(*44*, *45*) (*9*). Here we selected a *P. lactis* isolate (belonging to the wider *P. fluorescens* complex) from food and showed that it effectively competes with *Salmonella* in multiple conditions and environments and showed that the mechanisms driving its competitive fitness are environment dependent.

We first tested the potential of the selected strain *in vitro* and *in planta*. Interestingly we showed that the selected strain exhibited a competitive advantage over *Salmonella* in the plant environment using three distinct application timings. Notably, even when *Salmonella* was pre-established for 48 hours prior to *Pseudomonas* introduction, *Pseudomonas* could actively colonise the plant model and *Salmonella* numbers did not increase and remained stable over time.

Although multiple mechanisms of competition and suppressive activity employed by *Pseudomonas* strains have been characterised in the past (*8*, *46*, *47*), here we showed that there is no universal mechanism employed to compete against *Salmonella* in the different environments tested. Instead, our selected *Pseudomonas* strain (BA_HP_) underwent significant metabolic shifts which are likely to determine the mechanisms used to control *Salmonella* in an environment specific manner. Of note is the fact that well characterised competition systems such as secretion systems or genes producing known secondary metabolites were either not differentially expressed in our transcriptomic data, indicating that the mechanisms responsible for activity in the different environments are likely novel or a combination of mechanisms promoted by metabolic adaptability. For example, genes associated with pyoverdine production (*pvdA*, *pvdQ*), iron uptake (*fpvA*), type III secretion system (*vgrA*) were significantly downregulated *in vitro* and were not differentially expressed *in planta*. Indeed, BA_HP_ did not produce detectable siderophores on a Chrome Azurol S assay in the presence of *Salmonella* (data not shown).

We hypothesise that the upregulated genes in both environments likely contribute to metabolic reprogramming that enhances resource acquisition, secondary metabolite production and stress adaptation in response to *Salmonella* in an environment-specific way.

*In vitro*, despite the downregulation of pyoverdine production, we observed strong upregulation of genes encoding for the bacterioferritins *bfr_1* and *2*. These genes are involved in iron storage and regulation, indicating *Pseudomonas* sequesters iron using a different mechanism (to siderophore production) limiting its availability to *Salmonella,* and potentially affecting its virulence and growth, while maximising intracellular iron which could potentially be used in other pathways important for competition. Genes involved in pentose uptake (*araF*, *araG*, *rbsC*) and regulation of sugar catabolism (*araC*) were also significantly upregulated, despite glucose being the sole carbon source in the media. This suggests efficient utilisation of specific sugars, potentially depriving *Salmonella* of these resources. Upregulation of genes involved in pyrimidine degradation (*dht*, *preT* and *preA*) indicate a potential adaptation under stress via nutrient recycling in the presence of *Salmonella*. Several genes within the benzoate degradation and tryptophan metabolism pathways were upregulated. These pathways are responsible for the production of secondary metabolite precursors such as anthranilate and catechol (*48*), potentially facilitating the production of antimicrobial compounds against *Salmonella*. Observed upregulation of degradation of aromatic compounds may be in response to secondary metabolites produced by *Salmonella*. Further investigation is required to test this hypothesis and to investigate this interaction from the *Salmonella* perspective.

*In planta*, our data suggests a strategic adaptation to the plant environment under microbial competition. Specifically, upregulation of genes involved in the synthesis of the Cbb3-type cytochrome c oxidase subunit (*ccoP*1 and *ccoN*1_2) indicates adaptation to low-oxygen conditions in plant tissues, potentially giving *Pseudomonas* a competitive advantage over *Salmonella* growing in the same niche. We also observed upregulation of genes potentially involved in the detoxification of plant-derived reactive compounds (*nemR, nemA, yqhD*), enabling it to persist and effectively compete against *Salmonella* for niche. Upregulation of *adhB_*2 (alcohol dehydrogenase) and *ompW* (porin responsible for nutrient uptake) could indicate adaptation to using plant-derived nutrients (eg. alcohols) more effectively through outer membrane adaptation. Upregulation of peptidoglycan biosynthesis may be involved in membrane reinforcement to help *Pseudomonas* resist plant defences and effectively compete against *Salmonella*. Increased oxidative phosphorylation indicates increased energy production that could give *Pseudomonas* a competitive advantage over *Salmonella* in this setting.

There was a high variability observed between biological replicates in the plant environment, compared to extremely consistent findings across the *in vitro* samples. This variability likely reflects both technical challenges associated with RNA extraction and sequencing from complex plant tissues, and inherent biological heterogeneity—since not all extracted cells are expected to colonise similar niches or interact directly with *Salmonella* in the plant environment. This increased noise in our dataset effectively makes it more difficult to confidently detect genes with subtle changes in expression and potentially causing us to miss biologically relevant signals in this environment.

Despite this, our pathway enrichment analysis including the two more highly correlated pairs of samples identified multiple genes within the same pathways as being altered in expression in a consistent manner. This pathway-level convergence provides additional confidence in our findings, even in the context of higher noise. Future work will focus on functionally validating these genes and pathways and employing single-cell RNA sequencing, which will allow us to resolve cellular heterogeneity and improve detection sensitivity, thereby overcoming some of the limitations encountered with bulk RNA-seq approaches.

While our findings focus on a specific *Pseudomonas fluorescence* strain, the well-documented metabolic plasticity and ecological adaptability across the *Pseudomonas* genus (*49*) suggest that similar environment-driven responses are likely to be a common feature among other members of this group. Understanding which environmental triggers *Pseudomonas* senses to switch gene regulation in the different environments, is key to detangle community dynamics. This will both pave the way to advance our understanding on interspecies interactions in complex mixed-species communities and unveil what mechanisms are key for a rational application of these factors towards community engineering.

In conclusion, our findings demonstrate the robust biocontrol capabilities of *Pseudomonas*, suggesting a complex adaptive response to competition in different environmental settings. As a ubiquitous member of food systems and plant environments, its natural prevalence and ability to colonise plants enhances ecological compatibility and survival, key factors strengthening its appeal as a biocontrol agent.

---

**Figure S1:**
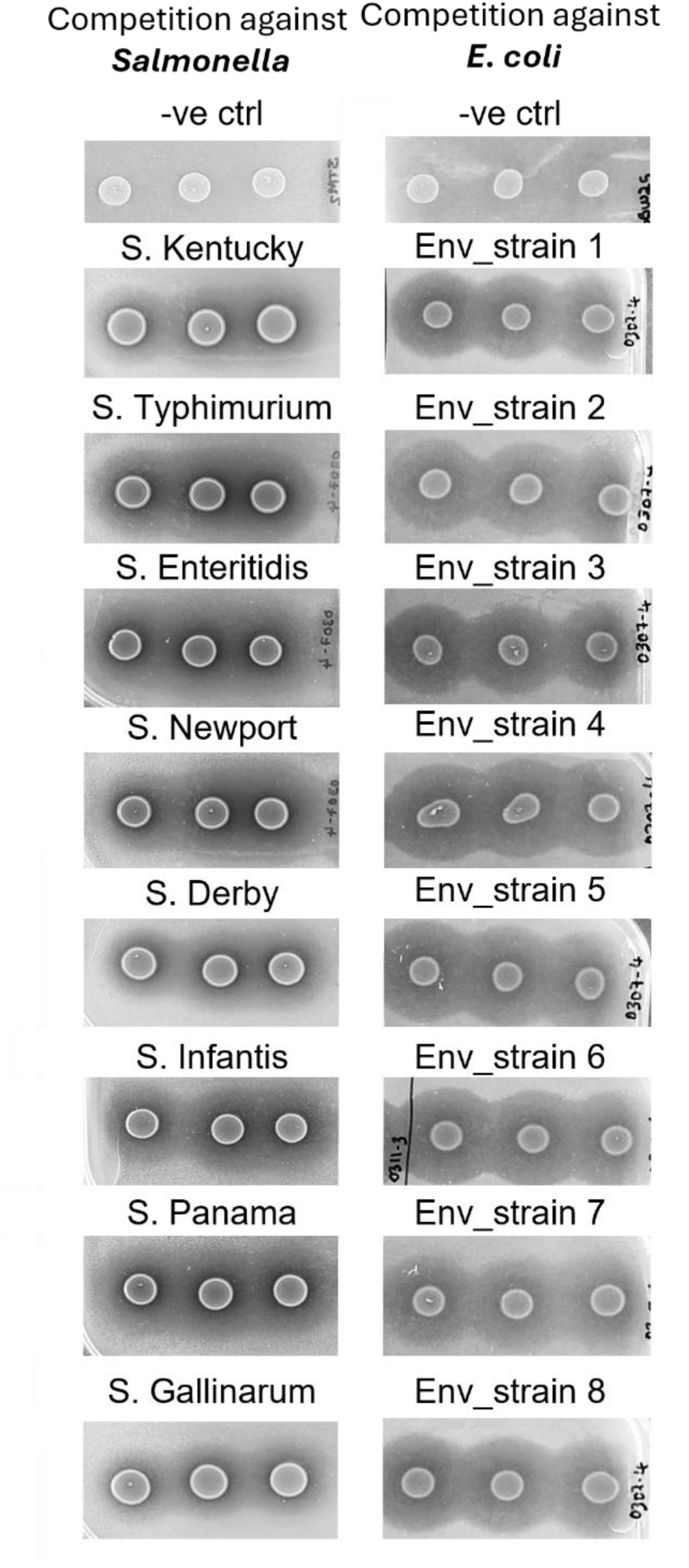
Competition assay of BA_HP_ against various Salmonella serovars and E. coli environmental isolates using an overlay assay on agar. Plates were incubated for 48 hours. Clearing zones indicate growth inhibition of the tested strains by BA_HP_.

**Figure S2:**
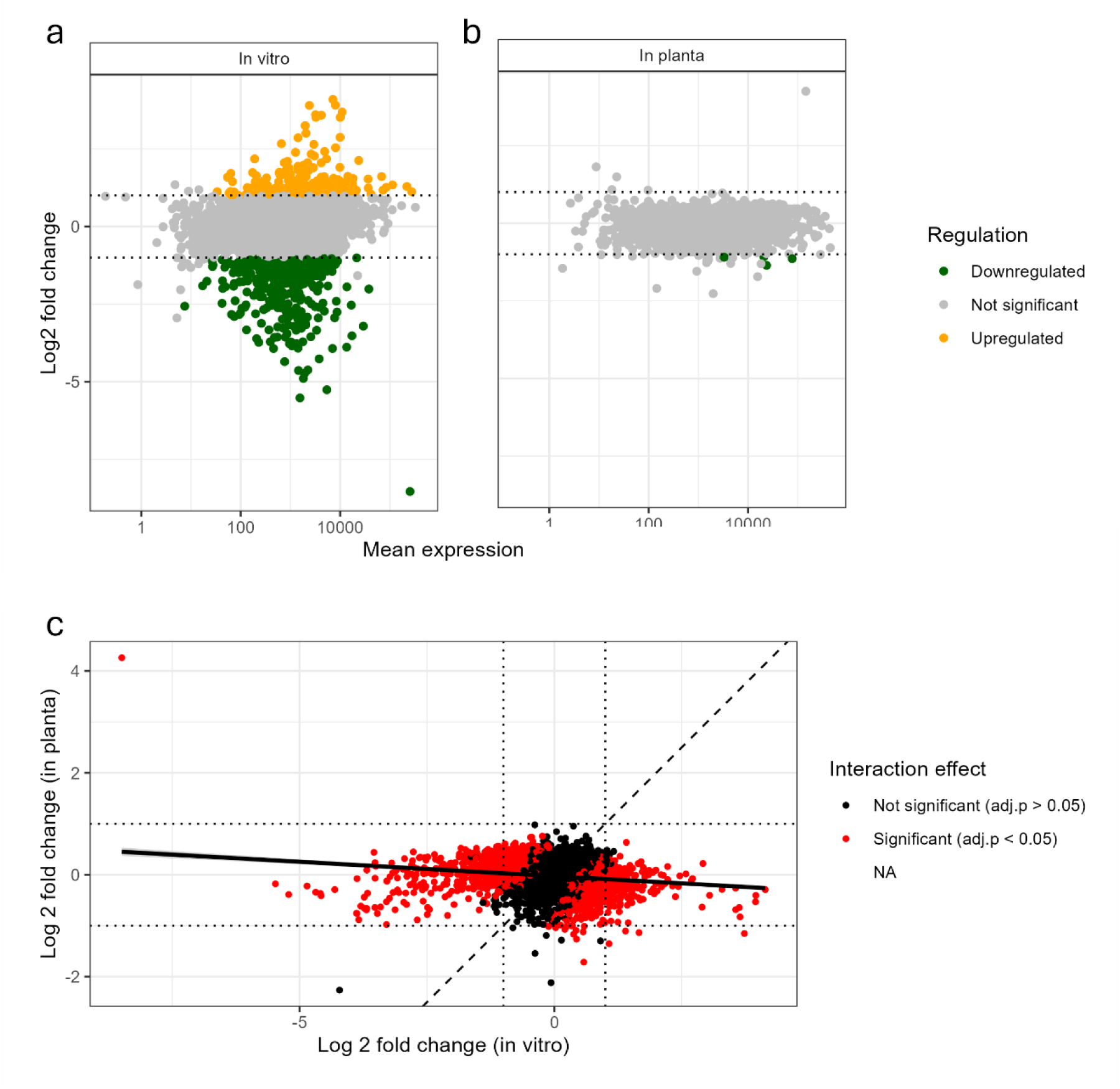
RNA sequencing results summary including all three in planta replicates. **a-b**: MA plots (log2 fold change vs. mean expression) showing the gene expression changes in response to Salmonella in vitro (a) and in planta (b). Changes in expression was only considered significant when p<0.05 and LogFC ≥ 1/≤ −1. Green dots represent downregulated genes, while orange dots represent upregulated genes. Grey dots are not significant. **c**. Scatter plot showing the relationship between log2 fold changes in gene expression under in vitro and in planta conditions. Each point represents a gene, with its log2 fold change in vitro on the x-axis and in planta on the y-axis. The solid black line indicates the best-fit linear regression, showing the overall trend. The dotted black line represents the identity line (y = x), where genes would fall if their expression changed equally in both conditions. Red points highlight genes with a statistically significant deviation from the expected relationship.

**Figure S3:**
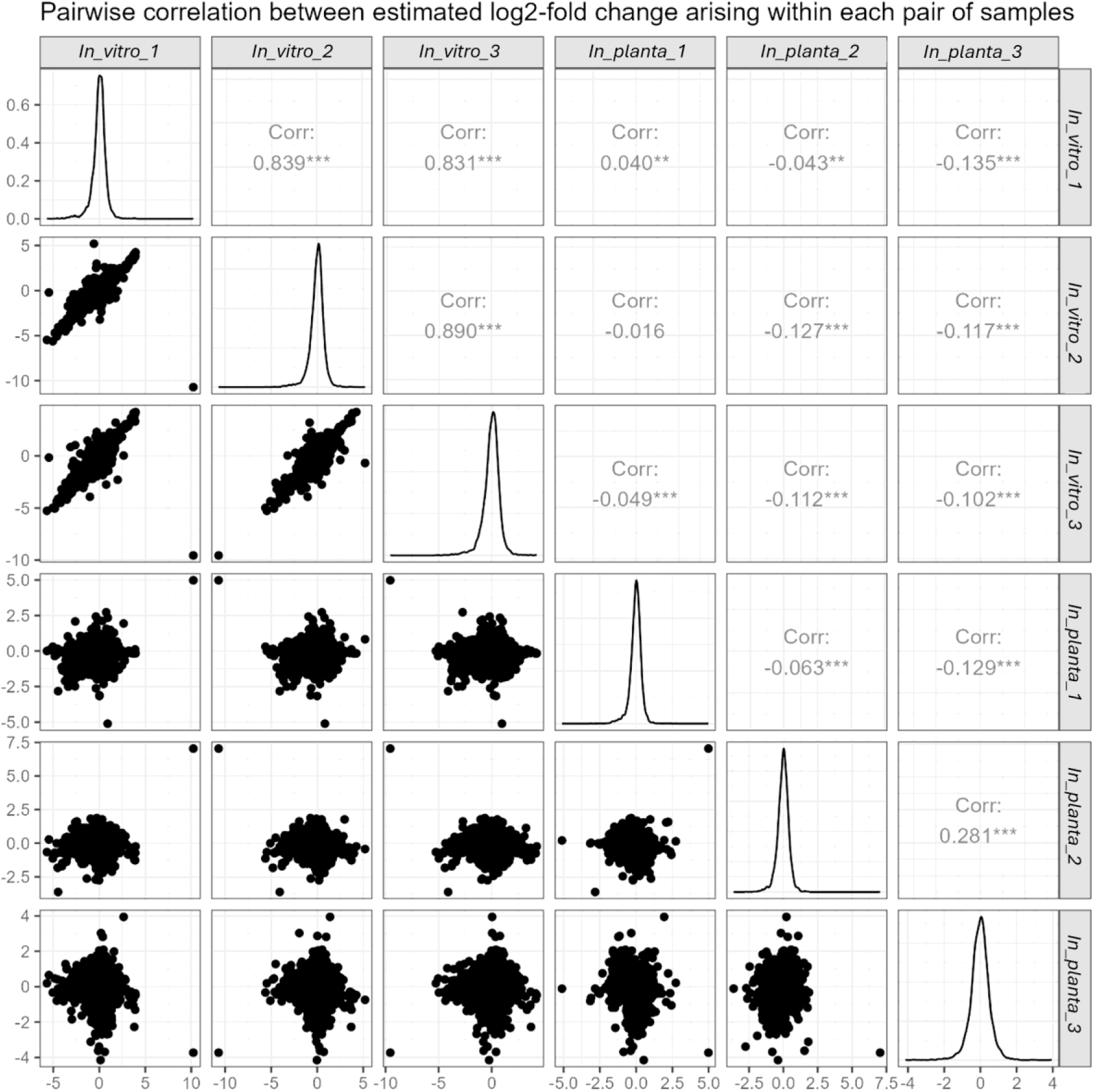
Scatter plots describing pairwise correlations of log2-fold changes for each sample pair, including all three replicates for both in vitro and in planta environments. Pearson correlation coefficients are reported for each pair, quantifying inter-sample concordance;

